# Graph-Based Classification of Penetrating Cortical Arteries and Veins in Whole-Mouse Vasculature Datasets

**DOI:** 10.64898/2026.10.15.758745

**Authors:** Sofia Farina, Letizia Greco, Giorgia Barra, Maxime Boyer, Nicolas Renier, Franca Schmid

## Abstract

Advances in whole-brain imaging have produced large-scale vascular datasets, yet most lack artery–vein annotations. However, this is essential for quantitative analysis of whole brain vascular structure, blood flow simulation and translational cerebrovascular research. For unlabelled datasets, manual labelling is infeasible at network scale, and surface-vessel tracing is hampered by reconstruction artefacts, limiting the insights that can be gained from these datasets.

Here, we focus on penetrating vessel trees, i.e. descending arteries (A) and ascending veins (V). A supervised Random Forest classifier is trained on graph-derived morphological and topological tree features, including persistent homology. Training uses datasets from two imaging modalities: Light sheet (LS) microscopy with arterial staining and two-photon (TP) micrscopy with manual labelling in smaller samples.

Classification reached 96% balanced accuracy on independent TP dataset and 93% on LS datasets. Topological features alone achieved competitive accuracy without relying on diameter measurements, demonstrating cross-modality portability. Arteries exhibit larger diameters and longer inter-bifurcation segments, likely facilitating their functional task of directed blood delivery. The veins show greater branching complexity, allowing low-pressure drainage. These signatures are stable across modalities, supporting their physological relevance.

The framework enables A-V label transfer to unlabelled whole-brain TP graph datasets, validated through the V:A ratio as a function of middle cerebral artery (MCA) distance. Operating on graph representations derived from any volumetric imaging modality and requiring no retraining, the approach is applicable to LS, TP, and likely other imaging platforms, providing a scalable foundation for A-V quantification, vascular topological analysis, and haemodynamic modelling.

## I. Introduction

Recent advances in high-resolution imaging techniques have enabled whole-brain vascular acquisitions in the mouse ranging from capillaries to large surface vessels [1]–[5]. These datasets have opened new opportunities for quantitative analysis of cerebrovascular organization across the entire brain. Moreover, the graph-base representation of vascular networks [5], [6] is essential for modelling cerebral blood flow and different processes essential for healthy brain function [7], [8].

Despite this progress, current whole-brain vascular datasets often suffer from important limitations. In particular, there is a trade-off between accurately resolving capillary-level features and faithfully representing larger surface vessels. This compromise can arise from multiple sources, including sample preparation, staining protocols, image acquisition, segmentation accuracy, and errors introduced during graph construction [9]. As a result, generating accurate and reliable vascular graphs suitable for downstream analysis is a persisting challenge in cerebrovascular research.

A critical requirement for physiologically meaningful hemodynamic modeling, vascular topological characterization, and clinical or translational interpretation of cerebrovascular networks is the annotation of vessel classes, in particular whether a branch is a capillary, an artery or a vein. Arterial and venous vessels play distinct physiological roles: arteries deliver oxygenated blood to the tissue and are responsible for blood flow regulation, while veins drain deoxygenated blood. Most neurological disorders have specific impacts on the structure and organization of veins and arteries. The specificity of arteriovenous alterations in each diseased state highlight the critical importance of accurate artery–vein labeling for understanding perfusion and their impact on neuronal functions. Moreover, most publicly available whole-brain vascular datasets lack artery–vein annotations [1], [2], [10], [11], severely restricting their applicability for cerebrovascular research.

Recent work has partially addressed this limitation by using immunostaining protocols targeting smooth muscle cells (SMC) [5], which are predominantly present in arterial vessel walls. Applied to *ex vivo* preparations, this approach enables the acquisition of dual-channel images consisting of an endothelial and a SMC channel, allowing the construction of labelled vascular graphs. However, targeted SMC staining alone does not ensure a straightforward identification of arterial vessels, as regional variations in signal intensity and the presence of signal in venous segments require manually tuned algorithms for reliable vessel classification. In addition to these challenges, for the vast majority of existing datasets such staining is generally unavailable and infeasible for most techniques.

The most intuitive approach to artery–vein labeling in unlabeled datasets would be to trace vessels from the major trunks at the brain surface into the capillary bed. In practice, however, this is not feasible: at the pial surface, arteries and veins frequently run alongside or cross one another, making reliable reconstruction and tracing difficult. To address this challenge, we focus on penetrating vascular trees, which originate from surface vessels and extend into the parenchyma as descending arteries (A) and ascending veins (V). Prior studies and expert anatomical knowledge have shown that A and V penetrating trees exhibit distinct topological and morphological characteristics in the cortex that allow manual differentiation by experts at the local scale [12]. However, manual labeling is impractical at the whole-brain level due to the size, heterogeneity, and complexity of the vascular network and automated A-V discrimination at the graph level remains largely unexplored.

In this work, we aim to overcome these limitations by developing a computational framework for A-V labeling of vascular graph. Since A and V differences are best characterised in the cortical vasculature [10], [12], we restricted the analysis to the isocortex in whole-brain datasets, as defined by the Allen Brain Atlas annotation. The main contributions of this work are threefold:

1. We propose a novel algorithm for extracting penetrating vascular trees from whole-brain graph representations of the mouse vasculature. The method robustly identifies penetrating vessels emerging from the cortical surface.
2. We characterise the morphological differences between A and V using two complementary datasets: lightsheet microscopy (LS) data with SMC staining, and partially labelled two-photon (TP) microscopy data. In particular, we leverage persistent homology through a Topological Morphology Descriptor (TMD) [13], [14], here applied for the first time to cortical penetrating trees as a topological fingerprint of artery–vein branching patterns. Building on this characterisation, we introduce an automated A-V classification framework based on graph-derived geometrical and topological features combined with supervised machine learning.
3. We demonstrate that our framework enables reliable label transfer to unlabelled TP whole-brain vascular dataset in the cortex, without requiring SMC staining at inference time. To validate this transfer, we compare the predicted middle cerebral artery (MCA) territory distribution against labelled LS reference datasets, demon-strating the potential of our approach for large-scale cerebrovascular analysis and haemodynamic modelling.

## II. Methods

### A. Data Acquisition and Graph Construction

We consider several datasets from two independent laboratories. The first group comprises three microvascular networks and one whole-brain vascular dataset, all acquired with TP microscopy. The microvascular networks (MVN1–MVN3) (Fig. 1 (a)–(c)) were extracted from the isocortical region and manually annotated with A and V labels. MVN1 and MVN2 were investigated in several previous studies [7], [15], [16] and acquired with a voxel resolution of 1.0 µm × 1.0 µm × 1.0 µm. The whole-brain dataset, referred to hereafter as TP-MVN3, was first published in [2] at a voxel resolution of 0.3 µm × 0.3 µm × 1.0 µm. A cortical cutout extracted from TP-MVN3, denoted MVN3, was subsequently manually labelled and used for topological analysis [10].

**Fig. 1.**
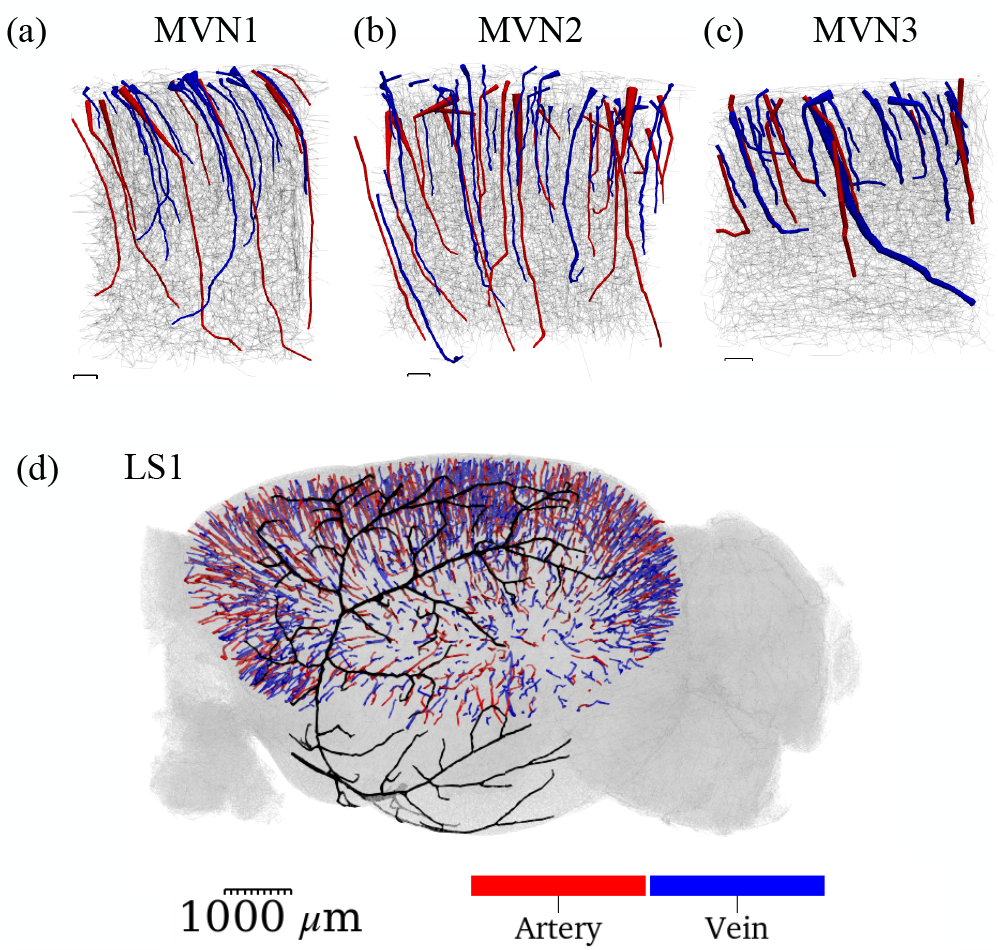
Graph representations of vascular datasets with labelled penetrating trees. (a) MVN1, (b) MVN2, and (c) MVN3: with manually labelled penetrating trees color-coded by vessel type (scale bar: 100 µm). (d) LS1 with penetrating vessel trees extracted from the isocortical region and color-coded by vessel type. Each tree was assigned a single label based on the SMC staining acquired at imaging time. The MCA was manually identified and highlighted in black (scale bar: 1000 µm).

The second group comprises four left-hemisphere vascular networks (LS1–LS4) [17] and acquired with LS microscopy at a resolution of 1.625 µm × 1.625 µm × 2.5 µm (Fig. 1(d)). These datesets are acquired following the protocol and an updated version of the reconstruction pipeline described in [5], with As and Vs identified by SMC staining and a manually tuned identification algorithm. All datasets were provided in graph format, with vertices and edges extracted using each laboratory’s own image processing pipeline [2], [5].

These pipelines produced graph representations of the vascular network together with geometric and spatial measurements derived from the underlying microscopy images. In this representation, nodes correspond to vessel bifurcations or endpoints, and edges connect pairs of adjacent nodes where the corresponding vessel segments are described as a sequence of segments.

Each node was associated with its spatial coordinates and a distance-to-surface measurement, important for the penetrating tree algorithm extraction. For the LS1–LS4 datasets, the distance-to-surface values were given from the corresponding post-processing pipeline. For TP-MVN3 datasets, the distance-to-surface metric was computed using an in-house algorithm applied to the graph. Each edge was assigned a diameter (median diameter of the vessel segments) and a length (sum of segment lengths). This common representation enabled consistent geometrical and topological analyses across datasets while retaining measurements from the original image-processing pipelines.

In whole-brain datasets LS1–LS4, the isocortical region was identified using the Allen Mouse Brain Common Coordinate Framework (CCFv3), whose annotations were incorporated during graph construction (Fig. 1(d)).

Subsequently, penetrating vascular trees were extracted from LS1–LS4 and the right hemisphere of TP-MVN3 using Algorithm 1. For MVN1–MVN3, the penetrating trees were identified based on the existing the A/V-annotation. A penetrating tree refers to the main trunk of a penetrating vessel, tracing a path from the cortical surface to its deepest point (Fig. 1).

The penetrating tree extraction algorithm requires parameter tuning, including the definition of candidate surface roots based on their distance to the cortical surface, and the weights used to score the optimal path. To select the best parameter configuration and validate the algorithm, we applied it to TP-MVN3, using the manually annotated penetrating trees of MVN3 as a reference. Parameters were optimised to maximise the recovery of annotated trees and their edges, while minimising false-positive edges. The best configuration recovered 36 of 38 penetrating trees (94.74%), with an edge recall of 76.47% and a precision of 65.86%.

The algorithm was subsequently applied to the cortical regions of the LS1–LS4 datasets (Fig. 1). Due to differences in the imaging modality and the resulting graph metrics compared with the TP-MVN3, some parameters required manual adjustment. Furthermore, an additional selection step was necessary to recover large veins that were not captured by the initial extraction, but could be readily identified from the labelled edges in the graph. The number of penetrating trees in each dataset is reported in Table I.

**TABLE I.**
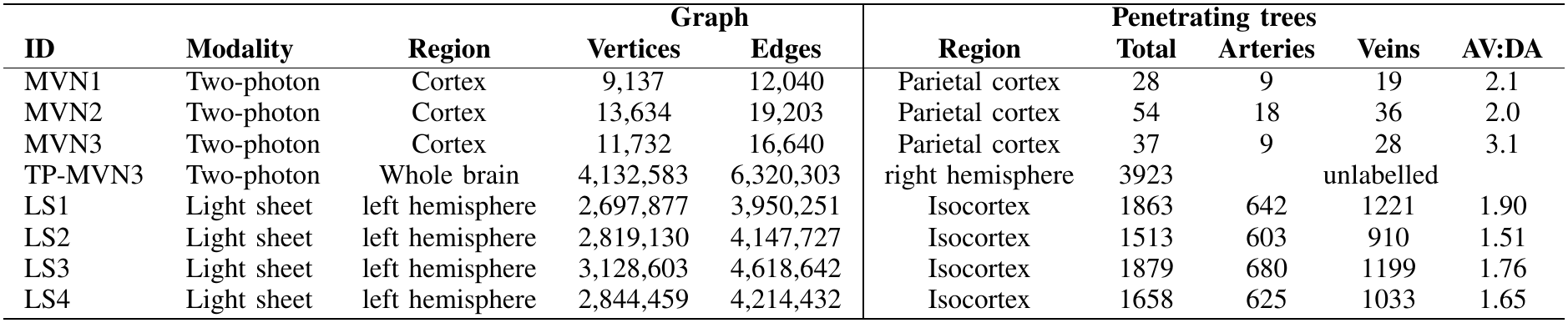
Dataset summary and penetrating tree statistics.

|  |  |  | Graph |  | Penetrating trees |  |  |  |  |
|---|---|---|---|---|---|---|---|---|---|
| **ID** | **Modality** | **Region** | **Vertices** | **Edges** | **Region** | **Total** | **Arteries** | **Veins** | **AV:DA** |
| MVN1 | Two-photon | Cortex | 9,137 | 12,040 | Parietal cortex | 28 | 9 | 19 | 2.1 |
| MVN2 | Two-photon | Cortex | 13,634 | 19,203 | Parietal cortex | 54 | 18 | 36 | 2.0 |
| MVN3 | Two-photon | Cortex | 11,732 | 16,640 | Parietal cortex | 37 | 9 | 28 | 3.1 |
| TP-MVN3 | Two-photon | Whole brain | 4,132,583 | 6,320,303 | right hemisphere | 3923 | unlabelled |  |  |
| LS1 | Light sheet | left hemisphere | 2,697,877 | 3,950,251 | Isocortex | 1863 | 642 | 1221 | 1.90 |
| LS2 | Light sheet | left hemisphere | 2,819,130 | 4,147,727 | Isocortex | 1513 | 603 | 910 | 1.51 |
| LS3 | Light sheet | left hemisphere | 3,128,603 | 4,618,642 | Isocortex | 1879 | 680 | 1199 | 1.76 |
| LS4 | Light sheet | left hemisphere | 2,844,459 | 4,214,432 | Isocortex | 1658 | 625 | 1033 | 1.65 |

### B. Ground Truth Label Validation

For LS1–LS4, each edge carries an artery, vein, or capillary label assigned during reconstruction. Artery labels are derived from the SMC staining, which generates a per-pixel intensity

#### Algorithm 1

Penetrating Tree Extraction from Vascular Graphs

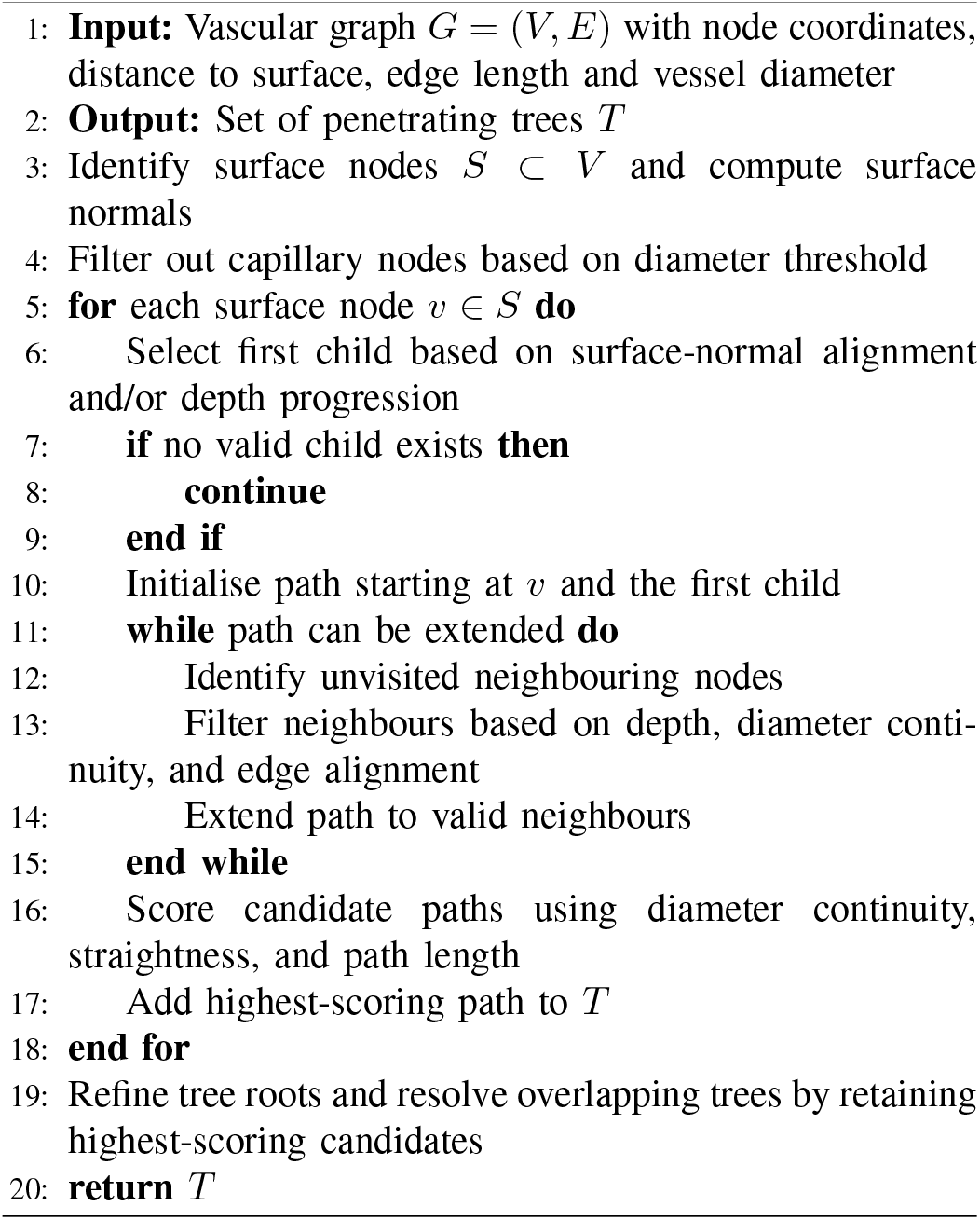

map. This map is processed using `ClearMap` [5] to assign A and V labels to graph edges based on signal intensity, vessel diameter thresholds, and subsequent propagation along the vascular network through graph connectivity.

Although this procedure provides an initial indication of vessel identity at the edge level, we observed that edge labels within penetrating trees were inconsistent in a few instances. In particular, vessels closer to the cortical surface tend to retain clearer artery or vein labels, whereas deeper segments are more frequently labelled as capillaries. Consequently, a single penetrating tree may contain a mixture of artery, vein, and capillary labels. In rare cases, a tree may contain both artery and vein edges, with or without capillary edges. This may reflect limitations in signal quality, graph connectivity, or `ClearMap` parameter choices during label propagation.

To obtain a definitive label (ground truth, GT) for each penetrating tree, we applied a majority-voting over artery and vein edges, excluding capillary-labelled edges. Trees composed entirely of capillary-labelled edges were assigned a vein label. In cases of a tie between A and V edges an edge-volume weighted vote was applied. (see the final penetrating tree label assigned for LS1 in Fig. 1).

To assess the quality of these labels, we examined the ratio of ascending venules to descending arteries (V:A), which in mice is known to lie between 2 and 3 [15], [18], [19]. MVN1–MVN3 satisfy this expectation. However, LS1–LS4 yield ratios between 1 and 2 (see Table I). This discrepancy might imply that the V:A ratio varies across brain regions or that some venular trees have been missed by the penetrating tree extraction algorithm.

To further validate the penetrating tree extraction algorithm and the `ClearMap`-A/V-labelling, we manually inspected several extracted graph-trees by overlaying them on the raw images and examining the SMC channel. Representative examples of an artery with a strong signal in the arterial channel and a vein with no arterial signal are shown in Fig. 2(a).

**Fig. 2.**
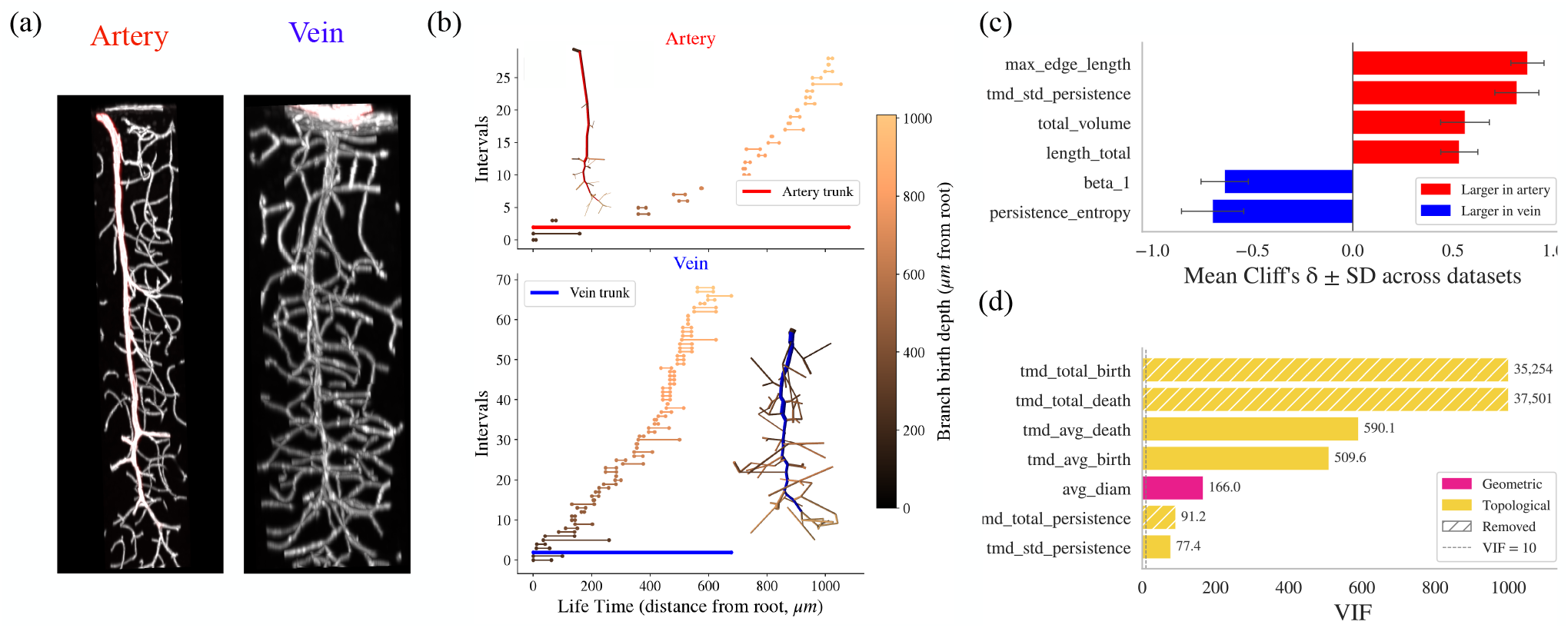
Label validation and feature extraction. (a) Representative penetrating artery and vein from dataset LS4. SMC staining channel in red (b) TMD barcode of an artery (top) and vein (bottom), with shared x-axis (Euclidean distance from root). The main trunk life-time interval is shown in red (artery) and blue (vein); branch intervals are colour-coded by birth value (distance from root at branching point). Inset: graph representation of the corresponding penetrating tree, colour-coded to match the TMD intervals. (c) Mean Cliff’s ***δ*** across all datasets for a representative subset of features. Red bars indicate selected features larger in arteries (***δ >* 0**); blue bars indicate features larger in veins (***δ <* 0**). Error bars show the standard deviation across datasets. (d) Variance Inflation Factor (VIF) for a representative subset of features. Yellow bars denote topological features; pink bars denote geometrical features. Hatched bars indicate features removed from the analysis due to extreme redundancy and negligible discriminability.

### C. Feature Computation

For each penetrating tree in the graph representation, we extract two types of features. Geometrical features (GF) quantify measurable physical or spatial properties of the vascular tree derived from the image and assigned to the graph, thus affected by the measurement technique and resolution. Topological features (TF) describe the connectivity, hierarchy, and branching pattern of the graph-tree. Thereby, providing a more intrinsic measurement that only requires the coordinates of the vertices points, and thus is less sensitive to measurement errors and to variability introduced by differences in acquisition or processing pipelines across datasets.

#### 1) Geometrical Features

The extracted GF have previously been reported relevant for A and V characterization [10], [12]. Each feature is computed at the tree level, resulting in one value per penetrating vascular tree, for a total of 19 features:

- **Diameter** [µm]: mean, median, standard deviation, minimum, and maximum edge diameter.
- **Total tree length** [µm]: sum of the lengths of all edges within the tree, representing its overall spatial extent.
- **Depth** [µm]: maximum distance from the cortical surface to the deepest node in the tree.
- **Total Volume** [µm]: volume of the penetrating tree.
- **Edge length** [µm]: mean, median, standard deviation, minimum, and maximum edge length.
- **Bifurcations:** total number of bifurcation vertices within the penetrating trees.
- **Number of children:** total number of child branches originating from the tree at each bifurcation vertex.
- **Branching angles:** mean, median, minimum, and maximum angles between parent and child segments, describing the local branching geometry.

#### 2) Topological Features

Further, we quantified the topological branching structure of each penetrating tree using a topological morphological descriptor (TMD) [13], [14]. In addition to the main penetrating tree, we considered the second-order child branches associated with each vertex of the main tree, thereby incorporating both the penetrating tree and its surrounding vessels into the topological analysis. Briefly, starting from the root of each tree, the Euclidean distance of each branching point is used to construct a persistence representation of the branching hierarchy. Each branch is represented by a (*birth, death*) pair, where the birth and death values define the corresponding interval in the resulting barcode. The resulting persistence barcode (Fig. 2(b)) captures the hierarchical organization of the vascular branches and provides a topological representation of the branching structure extending from the main penetrating tree into the surrounding capillary bed.

Following the feature construction described in [20], we extracted the following 20 tree-level TF:

- **Number of intervals:** total number of intervals in the persistence barcode.
- **Persistence:** total, mean, median, standard deviation, minimum, and maximum persistence, together with entropy and concentration measures.
- **Birth:** total, mean, median, standard deviation and maximum birth value.
- **Death:** total, mean, median, standard deviation and minimum death value.
- **First Betti number:** number of independent topological loops in the vascular tree.

### D. Supervised Artery–Vein Classification

#### 1) Random Forest

We classified penetrating trees as A or V using three complementary feature representations: (i) geometrical features (GF), (ii) topological features (TF), and (iii) a combination of geometrical and topological features (GTF). For each feature representation, we trained a Random Forest (RF) classifier using LS1, MVN1, and MVN3 as training datasets. Hyperparameters were optimised using five-fold cross-validation with a grid search over the following ranges: number of trees ∈ {100, 150, 200, 300}, maximum depth ∈ {2, 3, 5, 10}, minimum samples to split ∈ {2, 5, 10}, minimum samples per leaf ∈ {1, 2, 4}, features per split `∈ {sqrt, log2}, None`, and class weight `∈ {balanced, balanced subsample, None}`. The hyperparameter optimization and classification were performed independently for each feature sets: geometrical, topological and geometrical and topological. The selected hyperparameters are reported in Table II. The resulting classifiers were evaluated on four independent datasets: LS2, LS3, LS4, and MVN2 and validated on TP-MVN3.

**TABLE II.** Optimised hyperparameters of the Random Forest classifier for each feature representation, selected via five-fold cross-validation.

| **Hyperparameter** | **GF** | **TF** | **GTF** |
|---|---|---|---|
| max_depth | 10 | 10 | 10 |
| max_features | sqrt | sqrt | sqrt |
| min_samples_leaf | 1 | 2 | 2 |
| min_samples_split | 2 | 2 | 5 |
| n_estimators | 200 | 100 | 200 |
| Class_weight | balanced | balanced_subsample | balanced |

#### 2) Support Vector Machine

To benchmark the RF classifier, we also trained a Support Vector Machine (SVM) for each of the three feature representations. The same pipeline was applied: features were standardised to zero mean and unit variance, reduced via PCA retaining 80 % of the explained variance, and then passed to an SVM with a radial basis function (RBF) or linear kernel. Hyperparameters were optimised using five-fold stratified cross-validation scored by balanced accuracy, with a grid search over regularisation strength *C* ∈ {0.1, 1, 10, 100}, kernel width *γ* ∈ {`scale, auto`, 0.001, 0.01, 0.1} (RBF only), kernel type ∈ `{rbf, linear}`, and class weight ∈ `{balanced, None}`. The best model was refitted on the full training set and evaluated on the same four independent test datasets used for the RF. The selected hyperparameters are reported in Table III.

**TABLE III.** Optimised hyperparameters of the SVM classifier for each feature representation, selected via five-fold cross-validation.

| **Hyperparameter** | **GF** | **TF** | **GTF** |
|---|---|---|---|
| *C* | 10 | 1 | 1 |
| Kernel | rbf | rbf | rbf |
| *γ* | 0.01 | auto | auto |
| Class weight | balanced | balanced | balanced |

## III. Results

### A. Feature Analysis

We analyse all features extracted from the labelled datasets LS1–LS4 and MVN1–MVN3 to assess their ability to discriminate arteries from veins and to guide feature selection for the classifiers.

Fig. 2(c) shows the mean Cliff’s delta (*δ*) averaged across all seven datasets; positive values indicate a feature is larger in arteries, negative values in veins. Features are consistent across datasets from different anatomical regions, acquisition modalities, and sample sizes.

A and V show distinct topological and geometrical characteristics. Arteries have longer vessel segments, as reflected by a higher maximum edge length (`max_edge_length`), indicating that the longest branch in arterial trees extends over greater spatial scales than in veins. This is consistent with their higher standard deviation of persistence (`tmd_std_persistence`): a few dominant branches persist over long spatial scales whilst many minor ones disappear quickly (Fig. 2(b)). Arteries also have a greater total path length (`length_total`) and larger calibre (`total_volume`).

Veins, in contrast, exhibit more complex branching. The first Betti number (`beta_1`) is higher in veins, indicating more loop-like structures in the surrounding vessels when considering the first two generations of children. This is consistent with longer capillary loops close to arteries [10], meaning they require more edges to close. The entropy of the persistence (`persistence_entropy`) is also higher in veins, indicating a more uniform distribution of persistence across the branching structure, with no single branch dominating the structure. Overall this reflects a more branching structure compared to arteries.

We compute the Variance Inflation Factor (VIF) to identify and remove redundant features, retaining only those that contribute independent information to the classifier (Fig. 2(d)). Three features exhibit extreme redundancy with negligible discriminability (|*δ*| *<* 0.12): total birth, total death and total persistence. These three features are excluded from the classifier input.

Although `tmd_avg_birth` and `tmd_avg_death` also show high VIF values (510 and 590, respectively), they are retained because both exhibit large effect sizes (*δ* = 0.57 and *δ* = 0.65) and removing them would discard genuine discriminative information. The final feature set comprises 19 geometric features (GF), 17 topological features (TF), and 36 combined features (GTF = GF ∪ TF).

### B. Classification performance

Prior to supervised classification, we verified that the extracted features do not trivially separate arteries from veins. K-means clustering (*k* = 2) yielded a silhouette score of 0.366 and an Adjusted Rand Index of 0.321 against the known labels; hierarchical clustering achieved higher label agreement (ARI = 0.51) but poorer cluster compactness (silhouette = 0.281). Neither method consistently recovered the biological A-V distinction.

Table IV summarises classification performance across feature sets comparing RF and SVM. RF consistently outperforms SVM in both accuracy and specificity. On LS2–LS4, the best-performing feature set is the combined GTF, although GF alone also yields strong results. On MVN2, TF and GTF achieve higher accuracy than GF, suggesting that topological features are crucial for generalisation to a different imaging dataset. ROC-AUC values are consistently high across all configurations, confirming strong discriminative capacity regardless of the decision threshold. RF matches or exceeds SVM in ROC-AUC in all configurations, and GTF with RF achieves perfect separation (100%) on MVN2, further supporting its robustness to inter-dataset variability.

**TABLE IV.** Performance of the Random Forest (RF) and Support Vector Machine (SVM) classifiers for artery–vein classification using geometrical (GF), topological (TF), and combined (GTF) feature representations. Both models were trained on LS-1, MVN1, and MVN3.

| **Features** | **Classifier** | **Test set** | **Accuracy** | **F1** | **Sensitivity** | **Specificity** | **ROC-AUC** |
|---|---|---|---|---|---|---|---|
| GF | RF | LS2–LS4 | 92% | 93% | 93% | 90% | 97% |
|  |  | MVN2 | 94% | 96% | 92% | 100% | 99% |
|  | SVM | LS2–LS4 | 90% | 92% | 94% | 84% | 96% |
|  |  | MVN2 | 90% | 93% | 97% | 77% | 96% |
| TF | RF | LS2–LS4 | 89% | 91% | 90% | 88% | 95% |
|  |  | MVN2 | 96% | 97% | 94% | 100% | 99% |
|  | SVM | LS2–LS4 | 88% | 90% | 91% | 83% | 92% |
|  |  | MVN2 | 94% | 96% | 94% | 94% | 97% |
| GTF | RF | LS2–LS4 | 93% | 94% | 94% | 92% | 98% |
|  |  | MVN2 | 96% | 97% | 94% | 100% | 100% |
|  | SVM | LS2–LS4 | 92% | 93% | 95% | 86% | 96% |
|  |  | MVN2 | 93% | 94% | 94% | 89% | 96% |

The confusion matrices of the RF (Fig. 3 (a)) confirm these trends. On LS2–LS4, misclassifications are distributed across both classes for all three feature sets, with GTF producing the fewest total errors (357), followed by GF (409) and TF (537). On MVN2, errors are rare: the geometrical model misclassifies three veins and two arteries, whilst both the topological and combined models misclassify two veins and no arteries, yielding identical confusion matrices on this dataset. Across both classifiers, the combined feature representation, GTF, achieves the highest and most consistent performance. The improvement over geometrical features alone is modest in accuracy, but is reflected in a lower total error count on LS2–LS4. TF alone perform slightly below the other two representations on LS2–LS4, yet achieve 100 % specificity on MVN2 with the same confusion matrix as GTF, indicating that on this dataset the TF alone are sufficient to avoid any misclassification.

**Fig. 3.**
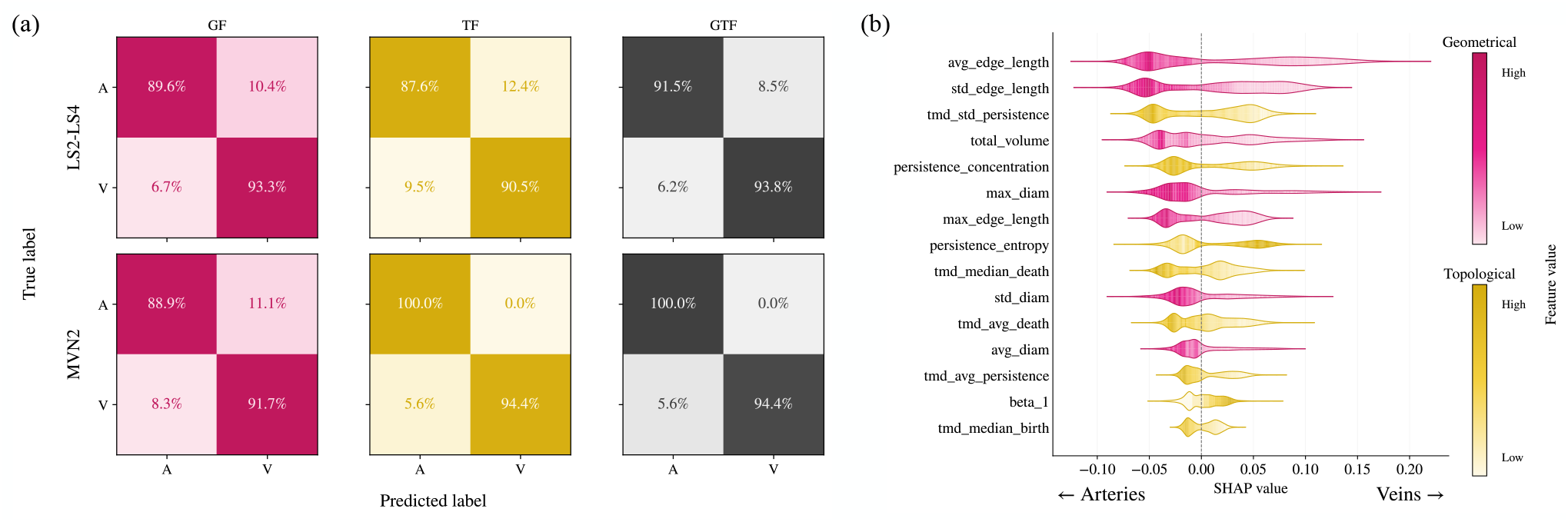
(a) Normalised confusion matrices for A–V classification on the independent test sets LS2–LS4 (top row) and MVN2 (bottom row), evaluated across three feature representations: geometrical (GF), topological (TF), and combined (GTF) (left to right). Values represent the percentage of vessels in each true class predicted as artery or vein. (b) SHAP feature importance for the combined GTF RF classifier, evaluated on the LS2– LS4 and MVN 2 test set. Features are ranked by mean absolute SHAP value. Pink violins denote geometrical features and yellow violins denote topological features with color intensity encodes the feature value (low to high). Negative SHAP values indicate a prediction towards arteries; positive values indicate a prediction towards veins.

To interpret the contribution of individual features, we performed SHAP (SHapley Additive exPlanations) analysis on the trained RF model for the GTF feature set, evaluated on the testing sets (LS2–LS4 and MVN2). Fig. 3(b) shows the SHAP feature importance for the combined model. Geometrical features are the most discriminative. Edge length features rank highest, with `avg_edge_length` and `std_edge_length` being the two most important features overall. High edge length values are associated with arteries, reflecting their longer inter-bifurcation segments and deeper cortical penetration, whilst veins exhibit shorter edges indicative of denser branching. The first topological feature, `tmd_std_persistence`, ranks third. High persistence variability is associated with arteries, suggesting greater heterogeneity in their topological structure. In contrast, `persistence_entropy` is higher for veins, consistent with a more uniform branching pattern. These findings are consistent with the feature analysis in Section III-A and known morphological differences between cortical arterial and venous trees [12].

### C. Analysis of misclassified trees

We investigate the misclassified trees in datasets LS2–LS4. The error rates are 8.1 % for GF, 10.6 % for TF, and 7.1 % for GTF. Combining geometrical and topological features therefore reduces misclassification compared to geometry alone, while topology alone produces the highest error rate.

We focus on trees that are misclassified by all three classifiers simultaneously (156 trees). For ground-truth arteries, a Kolmogorov–Smirnov analysis (Fig. 4(a)) identifies `tmd_std_persistence` and `max_edge_length` as the most discriminating features. The scatter plot of these two features (Fig. 4(b)) shows that misclassified arteries are structurally smaller than correctly classified ones. Their low persistence spread indicates a more homogeneous branching pattern, which is characteristic of veins rather than arteries.

**Fig. 4.**
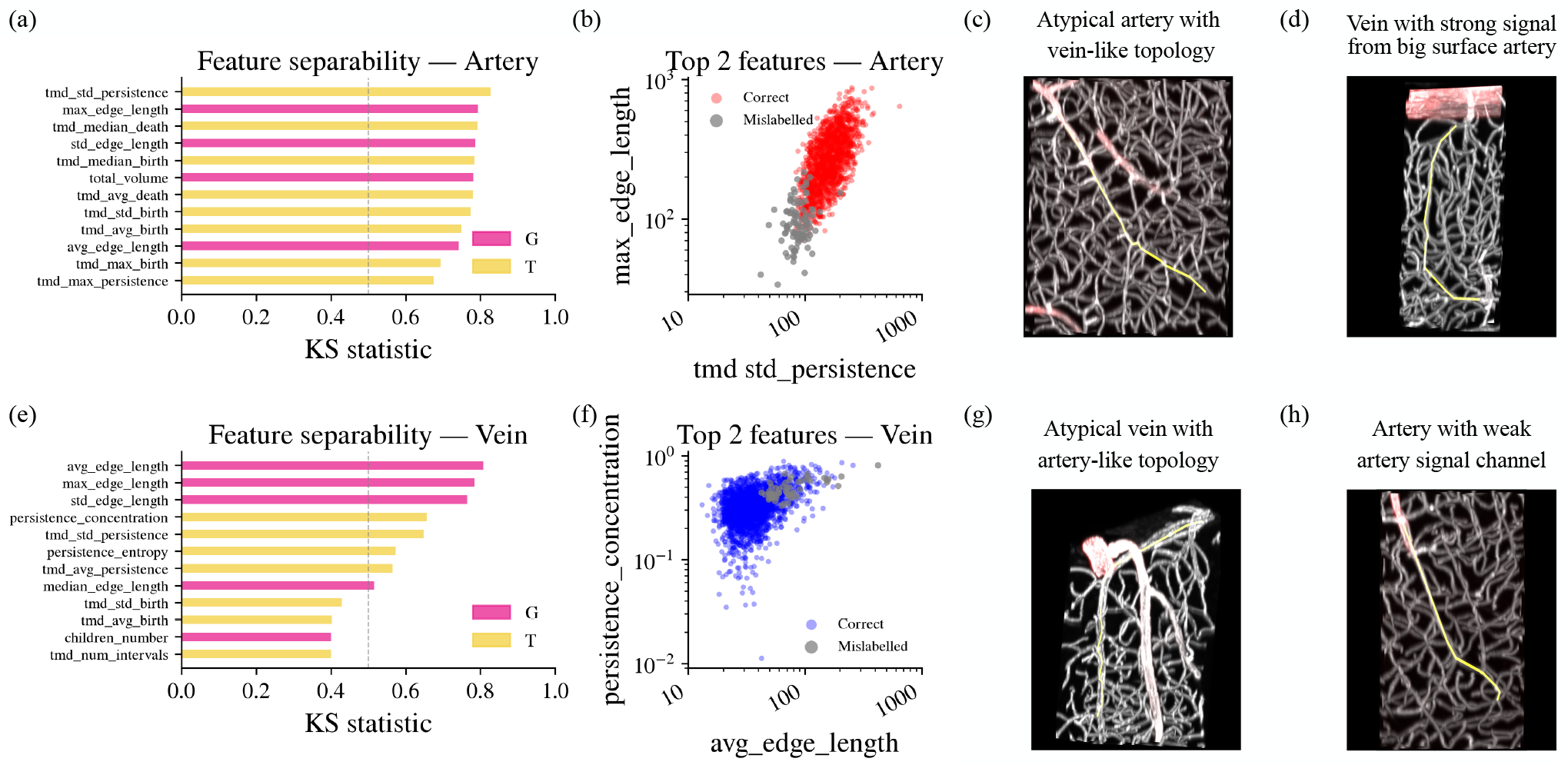
Analysis of trees misclassified by all three classifiers. (a) Kolmogorov–Smirnov (KS) statistic ranking of features based on their ability to separate correctly classified from misclassified ground-truth arteries; topological features (yellow) dominate the ranking. (b) Scatter plot of the two most discriminating features for arteries; misclassified trees (grey) cluster at lower values, indicating smaller and more homogeneous structures. (c)Microscopy image (grey scale) and artery signal channel (red scale) of a tree carrying an artery ground-truth label whose features led all three classifiers to predict vein. (d) A penetrating vein whose uppermost edges were incorrectly labelled as artery due to a strong artery signal present at the cortical surface; the label propagates into the graph but does not reflect the true vessel type. (e) KS statistic ranking for ground-truth veins; geometrical features (pink) rank highest. (f) Scatter plot of two of the most discriminating features for veins; misclassified trees show longer interbifurcation edges, a trait more typical of arteries. (g) A venous tree whose morphological features resemble arteries; the absence of artery signal confirms the vein-labelling and that the classifiers are incorrect. (h) A tree carrying a vein ground-truth label, likely due to a weak artery signal not captured during post-processing; the artery signal channel (red) indicates the vessel is an artery, consistent with the classifier predictions. G = geometrical; T = topological.

To better understand these errors, we manually inspected the 26 trees with a ground-truth artery label that all three classifiers misclassified as veins in LS4. For arteries, we identified three recurring cases. First, some arteries are genuinely atypical (around 27% cases): their morphological features closely resemble those of veins, and the classifiers make a plausible but incorrect decision. For example the tree in Fig. 4(c) shows many branches leading to more vein-typical persistence features. Second, around 66% of misclassified artery trees carry an incorrect ground-truth label due to edge-label propagation of the SMC signal. In some cases (35%), the extracted tree is a capillary or poorly captured penetrating artery: the tree-selection algorithm includes ambiguous segments, and the artery label propagates to them from an adjacent artery: a limitation of the tree-extraction step rather than of the labelling itself. In others (31%), the extracted tree is a genuine vein situated close to a surface artery with strong signal, which the `ClearMap` pipeline incorrectly propagates to the venous structure (Fig. 4(d)). In both cases, the classifiers correctly recognised the vein-like features of these trees; the error originates from upstream processing rather than from classification failure.

For ground-truth veins, the leading discriminating features are `max_edge_length, std_edge_length, avg_edge_length`, and `persistence_concentration` (Fig. 4 (e)). Misclassified veins tend to have longer inter-bifurcation edges and higher persistence concentration, indicating a dominant main branch rather than distributed ramification, a trait more typical of arteries (Fig. 4 (f)). Further, we investigated the 12 ground-truth veins mislabelled as arteries in LS4 by all three classifiers. First, some veins have morphological features following the expected classifier arterial pattern, around 25% cases(3/12). An example is shown in Fig. 4(g), where the tree has a long initial edge, large volume, and large diameter. Second, and around 33% cases (4/12), the ground-truth label itself appears erroneous: the artery signal channel shows clear staining, but the raw signal intensity was likely too weak at the time of labelling to be captured by the propagation algorithm. In these cases the classifiers correctly identify the tree as an artery, while the ground-truth records it as a vein (Fig. 4(h)). The remaining cases comprised capillary-like trees incorrectly selected by the extraction algorithm (2/12, 17%) and instances where the ground-truth label appears correct but the classifiers nevertheless failed (2/12, 17%).

In summary, the classifiers misclassified some trees whose geometrical and topological features more closely resemble those of the other vessel type. Visual inspection clarified the source of many errors. For arteries, the SMC channel was in some cases too weak to be captured by the ground-truth labelling algorithm, causing genuine arteries to be recorded as veins; the classifiers, however, correctly identified these vessels as arteries. For veins, the principal cortical venules [21] — large-diameter vessels — were systematically misclassified as arteries due to their atypical morphology, which is an outlier relative to the smaller penetrating veins. In both groups, a further source of error was the inclusion of poorly segmented trees or capillaries. After manual correction of 13 erroneous ground-truth labels in LS4, and excluding a separate set of 13 bad-tree or uncertain cases from the evaluation, the accuracy of all three classifiers increases by 0.79 percentage points, reaching 93.98%, 93.31%, and 95.44% for GF, TF, and GTF respectively. The uniform gain confirms that, in every corrected case, the classifiers had already predicted the true label, further supporting the validity of the pipeline.

### D. Application to an Unlabelled Whole-Brain Graph and Validation

To validate the framework on an unlabelled, unseen dataset, the trained RF classifiers were applied to TP-MVN3. The three feature representations (GF, TF, and GTF) were used independently to generate A-V predictions for all penetrating vascular trees extracted from the right hemisphere of TP-MVN3 (Table I and Fig. 5 (a)). Unlike the preceding analyses, penetrating trees here were extracted from the full left hemisphere of LS1-LS4 (Fig. 5 (b)).

**Fig. 5.**
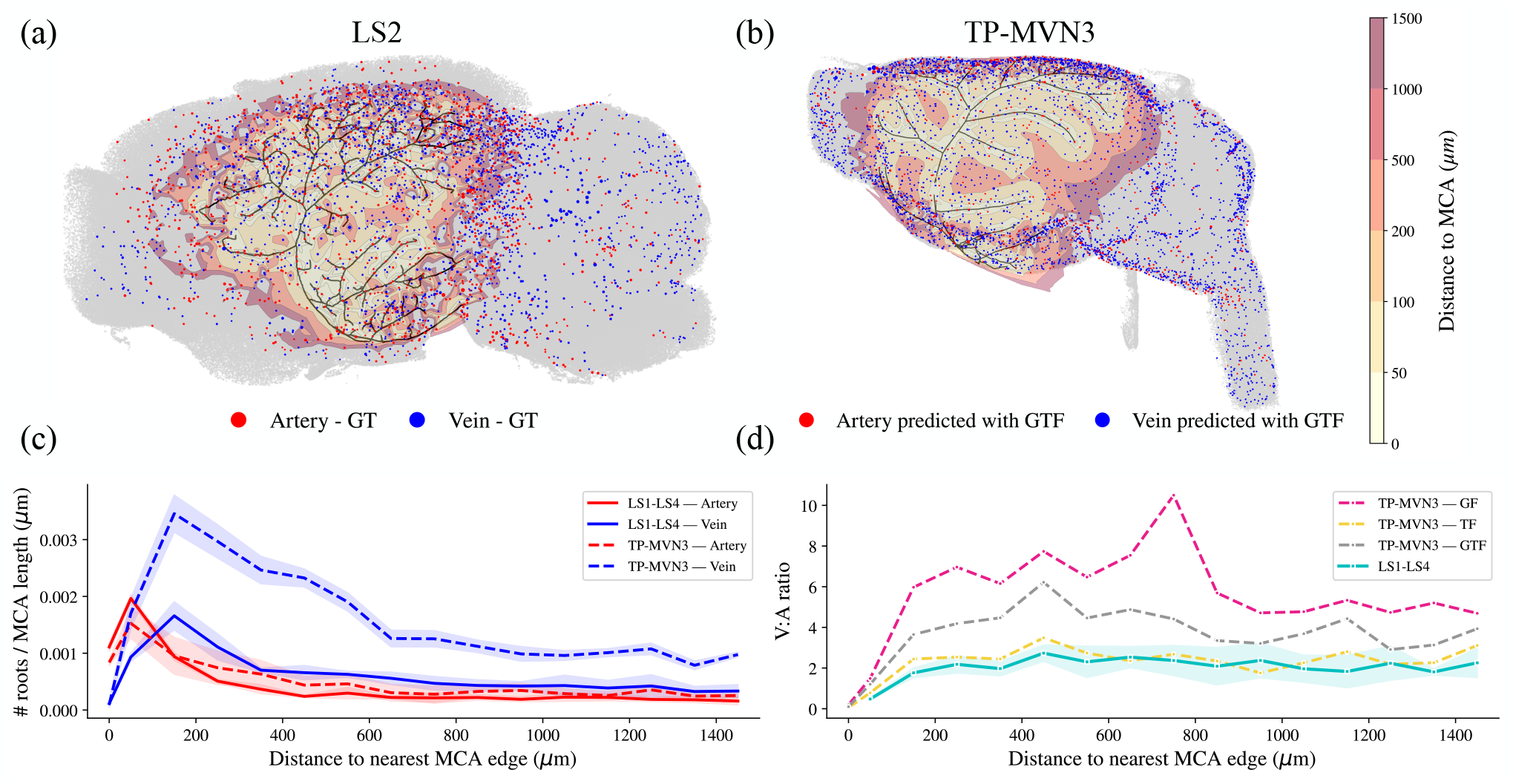
Azimuthal Lambert equal-area projection and spatial validation of artery–vein predictions. **(a)** LS-2 projected onto the left hemisphere, with the MCA highlighted in black. Penetrating arteries (red) and veins (blue) are color-coded according to ground truth. The highlighted zone indicates the region within 1500 µm of the MCA. **(b)** Same projection for TP-MVN3, with artery–vein labels predicted by the GTF classifier (arteries in red, veins in blue). **(c)** Penetrating tree roots, normalised by total MCA length, as a function of distance to the nearest MCA edge. LS1–LS4 arteries (red) and veins (blue) are shown alongside TP-MVN3 predictions averaged across the three classifiers (dashed lines); shaded areas indicate standard deviation across datasets for LS1–LS4 and across classifiers for TP-MVN3. **(d)** Vein-to-artery (V:A) ratio as a function of distance to the MCA. The LS1–LS4 mean and standard deviation are shown in cyan. Individual TP-MVN3 predictions are shown for GF (pink), TF (yellow), and GTF (grey).

The MCA was therefore used as a macroscopic anatomical landmark to assess the spatial distribution of predicted vessel classes. The MCA was manually identified in LS1– LS4 and TP-MVN3 (Fig. 5(a)-(b)). Both MCAs cover a similar anatomical region, yet structural differences are evident between datasets. TP-MVN3 contains 561 edges with relatively few surface vessel branches, whereas LS1–LS4 group averages 1839 ± 100 edges. This reflects differences in image acquisition modality, graph construction pipeline, and intra-subject anatomical variability. Despite these differences, the MCA provides a consistent reference point from which the local A-V distribution can be evaluated.

For validation, the analysis was restricted to vessels within 1500 µm of the MCA (Fig. 5(a)-(b)). As a spatial metric, we evaluated the number of penetrating tree root points normalised by MCA length as a function of distance to the MCA (Fig. 5(c)). On the MCA, arterial density is comparable between the two datasets (LS1–LS4: 0.0011; TP-MVN3: 0.0008), indicating consistent arterial recovery despite differences in MCA reconstruction. Vein density on the MCA is near zero, as expected, since penetrating veins do not originate directly from the MCA; the few detected likely reflect surface connectivity errors or misclassification. Moving away from the MCA, arterial density peaks in the proximal region and stabilises beyond 300 µm in both datasets. Vein density follows a similar trend, peaking around 200 µm before reaching a stable level. TP-MVN3 shows consistently higher vein density, suggesting that the optimised tree-extraction algorithm recovers a greater proportion of venous penetrating trees compared to LS1–LS4. This trend is further confirmed by the V:A ratio as a function of MCA distance for LS1–LS4 and for the three TP-MVN3 classifier predictions (Fig. 5(d)). Literature reports a V:A ratio of approximately 2–3 in the mouse cortical microvasculature [15], [18]. In LS1–LS4, the ratio stabilises beyond 300 µm, with a mean V:A of 1.99 (range 1.8–2.2 across subjects), consistent with reported cortical values. For TP-MVN3, all three classifiers show a similar initial trend, with arteries dominating near the MCA followed by a rapid increase in the V:A ratio. Beyond 300 µm from the MCA, the three classifiers diverge, with mean V:A ratios of 5.8 (GF), 2.5 (TF) and 4.0 (GTF), the GF excess reflecting a tendency to overclassify small penetrating arteries as veins. This bias peaks near 700 µm, where discordant trees, those labelled vein by GF but artery by both TF and GTF, are shorter (median length 624 vs 887 µm), shallower (median depth 427 vs 599 µm) and thinner (median max diameter 9.6 vs 12.0 µm) than the consensus artery population. Topological features mitigate this bias, as persistence-based descriptors are size-independent. Without ground-truth labels for TP-MVN3, a definitive classifier ranking cannot be established; however, the GF peak substantially exceeds the expected physiological range, raising concerns about its reliability. Furthermore, the V:A ratios reported in the literature are derived from small cortical samples [15], [18], and may not generalise to the full isocortical region analysed here. The expected V:A ratio at the hemisphere scale therefore remains uncertain, which limits the use of literature values as a validation benchmark.

In summary, the RF classifiers trained on labelled light-sheet data (LS1–LS4) generalise to an unlabelled two-photon dataset (TP-MVN3) without retraining. The predicted A-V distributions preserve the expected near-MCA artery dominance. The V:A ratio is higher in TP-MVN3 than in LS1– LS4, partly explained by the improved vein recovery of the calibrated tree-extraction algorithm. GF predictions show a further elevation, suggesting that geometric features alone overestimate venous prevalence across datasets. Balancing geometric with topological features is therefore preferable to reduce sensitivity to inter-dataset variability.

## IV. Conclusion

We presented the first framework to automatically classify arteries and veins in whole mouse-brain vascular graphs derived from different imaging modalities, and without dedicated staining at acquisition time. The framework builds on an algorithm to extract penetrating trees from large-scale vascular graphs and uses geometrical and topological features to train a classifier. Particularly, we apply persistent homology to brain vascular graphs extending topological data analysis beyond its prior use in neuronal morphology [13], [14] and vasculature [22], [23]. We validated the framework on an unlabelled two-photon dataset, where predicted labels reproduced the expected spatial organisation of arteries and veins around the MCA territory.

Both SVM and RF classifiers achieved high accuracy. The combined feature set (GTF) performed best, reaching 96% balanced accuracy on the TP dataset and 93% on the LS datasets. The accuracy gain from combining geometric and topological features over using either type alone was modest, which has a practical implication: geometrical features depend on imaging resolution and pipeline calibration, whereas topological features require only bifurcation coordinates and are therefore more portable. The fact that topological features alone achieve competitive accuracy suggests that robust classification can be obtained even without reliable diameter measurements. Manual inspection of misclassified trees further suggests that a substantial fraction of errors reflect label noise rather than classifier failure, so the reported accuracy is likely a conservative estimate of true performance. Based on overall performance and interpretability, we recommend the RF classifier for future applications.

SHAP analysis identified inter-bifurcation segment length as the most discriminating feature across all feature sets. This finding is functionally interpretable. Arteries exhibit longer and straighter segments, consistent with their role in directed, pressure-driven delivery of blood to specific cortical territories. Veins show greater branching complexity and are surrounded by denser capillary beds, supporting their role as distributed drainage pathways that collect blood across arterial territories and over cortical depth. These structural differences are consistent across imaging modalities and datasets, suggesting they reflect genuine biological organisation.

The ability to retrospectively assign A–V labels expands the potential of existing whole-brain vascular datasets. The present framework enables retrospective A–V labelling, facilitating systematic investigation of vascular organisation and blood-flow dynamics beyond small cortical volumes [7]. It also opens the door to quantifying penetrating artery density across subjects and strains: recent work suggests that differences in arterial density between mouse strains may impact autoregulatory capacity and tissue vulnerability [8], and the present framework provides a principled foundation for such comparative studies at scale.

A persistent challenge is vasculature segmentation and centerline extraction, which affect both tree detection and feature measurement. Recent advances in vessel segmentation, driven by deep learning and improved imaging protocols [24], are promising in this regard and are likely to yield cleaner inputs that further improve classification in the future.

Several promising directions offer opportunities to further enhance the framework. Extending it to additional imaging modalities would broaden its applicability across experimental platforms, although labelled data across modalities are currently scarce [1], [3], [11]. Extension to other brain regions would further test its generalizability; however, marked differences in vascular topology require thorough validation beyond the isocortex. Taken together, we provide a versatile frame-work that can establish artery–vein classification as a routine step in whole-brain vascular analysis, with its potential further strengthened by increasingly comprehensive and high-quality vascular datasets. This will support systematic comparisons across disease models, experimental conditions, and imaging platforms.

## Acknowledgment

The authors thank Dr. Xiang Ji and Prof. David Kleinfeld for data access, Gabrièle Lienhard for her contribution to the light-sheets datasets, and Ana Barrio for converting TP-MVN3 to the common framework. SF thanks Sergio Serrano for discussions at ECMTB2026.

